# vRNA-vRNA interactions in influenza A virus HA vRNA packaging

**DOI:** 10.1101/2020.01.15.907295

**Authors:** Sho Miyamoto, Yukiko Muramoto, Keiko Shindo, Yoko Fujita, Takeshi Morikawa, Ryoma Tamura, Jamie L Gilmore, Masahiro Nakano, Takeshi Noda

**Affiliations:** Laboratory of Ultrastructural Virology, Institute for Frontier Life and Medical Sciences, Kyoto University, Kyoto, Japan; Department of Molecular Virology, Graduate School of Medicine, Kyoto University, Kyoto, Japan; Keihanshin Consortium for Fostering the Next Generation of Global Leaders in Research, Kyoto, Japan; Laboratory of Ultrastructural Virology, Graduate School of Biostudies, Kyoto University, Kyoto, Japan

**Author notes:** Corresponding author (TN).

## Abstract

The genome of the influenza A virus is composed of eight single-stranded negative-sense RNA segments (vRNAs). The eight different vRNAs are selectively packaged into progeny virions. This process likely involves specific interactions among vRNAs via segment-specific packaging signals located in the 3’ and 5’ terminal coding regions of vRNAs. To identify vRNA(s) that interact with hemagglutinin (HA) vRNA during genome packaging, we generated a mutant virus, HA 5m2, which possessed five silent mutations in the 5’ packaging signal region of HA vRNA. The HA 5m2 virus had a specific defect in HA vRNA incorporation, which reduced the viral replication efficiency. After serial passaging in cells, the virus acquired additional mutations in the 5’ terminal packaging signal regions of both HA and PB2 vRNAs. These mutations contributed to recovery of viral growth and packaging efficiency of HA vRNA. A direct RNA-RNA interaction between the 5’ ends of HA and PB2 vRNAs was confirmed *in vitro*. Our results indicate that direct interactions of HA vRNA with PB2 vRNA via their packaging signal regions are important for selective genome packaging and enhance our knowledge on the emergence of pandemic influenza viruses through genetic reassortment.

## Introduction

The genome of the influenza A virus consists of eight segmented, single-stranded, negative-sense viral RNAs (vRNAs). Each vRNA contains a central coding region in an antisense orientation. This region is flanked by segment-specific noncoding regions and common terminal promoter sequences. Each vRNA forms a helical, rod-shaped ribonucleoprotein complex (vRNP) that associates with multiple nucleoproteins (NPs) and with a heterotrimeric RNA-dependent RNA polymerase complex composed of PB2, PB1, and PA proteins. The vRNP is responsible for transcription and replication of constituent vRNA. Recent studies examining NP-vRNA interactions in the context of vRNPs have shown that the NPs bind to vRNA nucleotides non-uniformly without sequence specificity[1, 2], suggesting that some parts of the vRNAs are free of NPs, and can potentially form secondary or tertiary structures that protrude from the surface of rod-shaped vRNPs.

There is evidence to suggest that progeny virions selectively package each copy of the eight vRNAs. In virions, the eight different vRNPs are arranged in a specific ‘1+7’ pattern, where one vRNP is surrounded by the other seven vRNPs[3, 4]. The mechanism by which each copy of the eight vRNAs is selected from a large pool of vRNAs and non-viral RNAs in virus-infected cells, and how these vRNAs are organized into the specific ‘1+7’ arrangement, remains unclear. The segment-specific packaging signal sequences, located in the noncoding and terminal coding regions of both the 3’ and 5’ ends of each vRNA, likely ensure the integrity of genome packaging[5–10]. The terminal coding regions within these packaging signal sequences are thought to be involved in co-packaging of multiple vRNAs, and are referred to as bundling signals [11]. Mutations or deletions in bundling signal sequences reduce the packaging efficiency of several vRNAs. The impact of such mutations or deletions on packaging efficiency is hierarchical among the eight vRNAs[8, 12–19], suggesting that there are specific interactions among the vRNPs. The vRNPs form sub-bundles *en route* to the plasma membrane [20, 21], and vRNPs in virions are directly or indirectly interconnected with each other[22, 23]; these findings offer additional evidence for the existence of vRNA-vRNA interactions.

To further examine the likely involvement of vRNA-vRNA interactions in genome packaging, it is necessary to identify which vRNA segments interact with one another, and which regions of each vRNA segment participate in these interactions. The potential *in-vitro* interactions of various naked vRNA segments have previously been described in human H3N2 and avian H5N2 viruses [22, 24]. However, the various combinations of vRNA-vRNA interactions differ between the two viruses. It also remains unclear whether such *in-vitro* vRNA-vRNA interactions in the absence of NPs reflect interactions that may occur among vRNPs *in vivo*. Only some nucleotides that are important for vRNA-vRNA interactions have been identified in the context of virus replication in cells and co-packaging into virions[22, 25]. These studies suggest that the 5’ ends of M vRNA and the central coding region of PB1 vRNA are involved in interactions and co-packaging with NA vRNA, respectively. In addition, interactions between PB1 and NS vRNAs may also be necessary for efficient viral replication and genome packaging. However, the region in NS vRNA identified to interact with PB1 vRNA is not located in the region of the previously reported genome packaging signal [26]. Thus, the role of vRNA-vRNA interactions in genome packaging remains unclear.

In this study, we used serial passaging to identify which vRNA(s) interact with the hemagglutinin (HA) vRNA packaging signal in selective genome packaging. We first generated a mutant influenza virus (A/WSN/33) that possesses five silent mutations in the packaging signal of HA vRNA, which causes a specific defect in the incorporation of HA vRNA. Then, we serially passaged the mutant virus in cultured cells to restore efficient incorporation of HA vRNA, and identified mutations that had been newly introduced into the vRNAs. In addition, we examined the interactions of HA vRNA with potential partner vRNAs *in vitro*, and assessed the importance of these vRNA-vRNA interactions in the packaging signal regions specific for HA vRNA incorporation and viral replication.

## Results

### Generation of mutant viruses possessing silent mutations in the packaging signal of HA vRNA

We first disrupted the packaging signal sequence of HA vRNA which potentially interacts with other vRNAs during selective genome packaging. For this, we used reverse genetics to generate a series of mutant viruses that possessed five silent mutations in either the 3’ or 5’ packaging signal region of HA vRNA without any amino acid mutations (Fig 1A). The respective viral titers were examined by plaque assays (Fig 1B). Eight out of nine mutant viruses replicated at a level similar to that of the wild-type virus, showing titers of approximately 3.8×10^8^ PFU/ml; however, an HA 5m2 virus, possessing five silent mutations at nucleotides 1664 to 1676 in HA vRNA, exhibited an approximately 87% reduced growth rate compared to the wild-type virus. Sequence analysis confirmed that no unexpected mutations occurred in any of the eight vRNAs of the HA 5m2 virus. These results indicate that nucleotides 1664 to 1674 in the packaging signal region of HA vRNA are necessary for efficient virus growth and are likely involved in HA vRNA packaging, which in agreement with results reported by a previous study[14].

**Fig 1.**
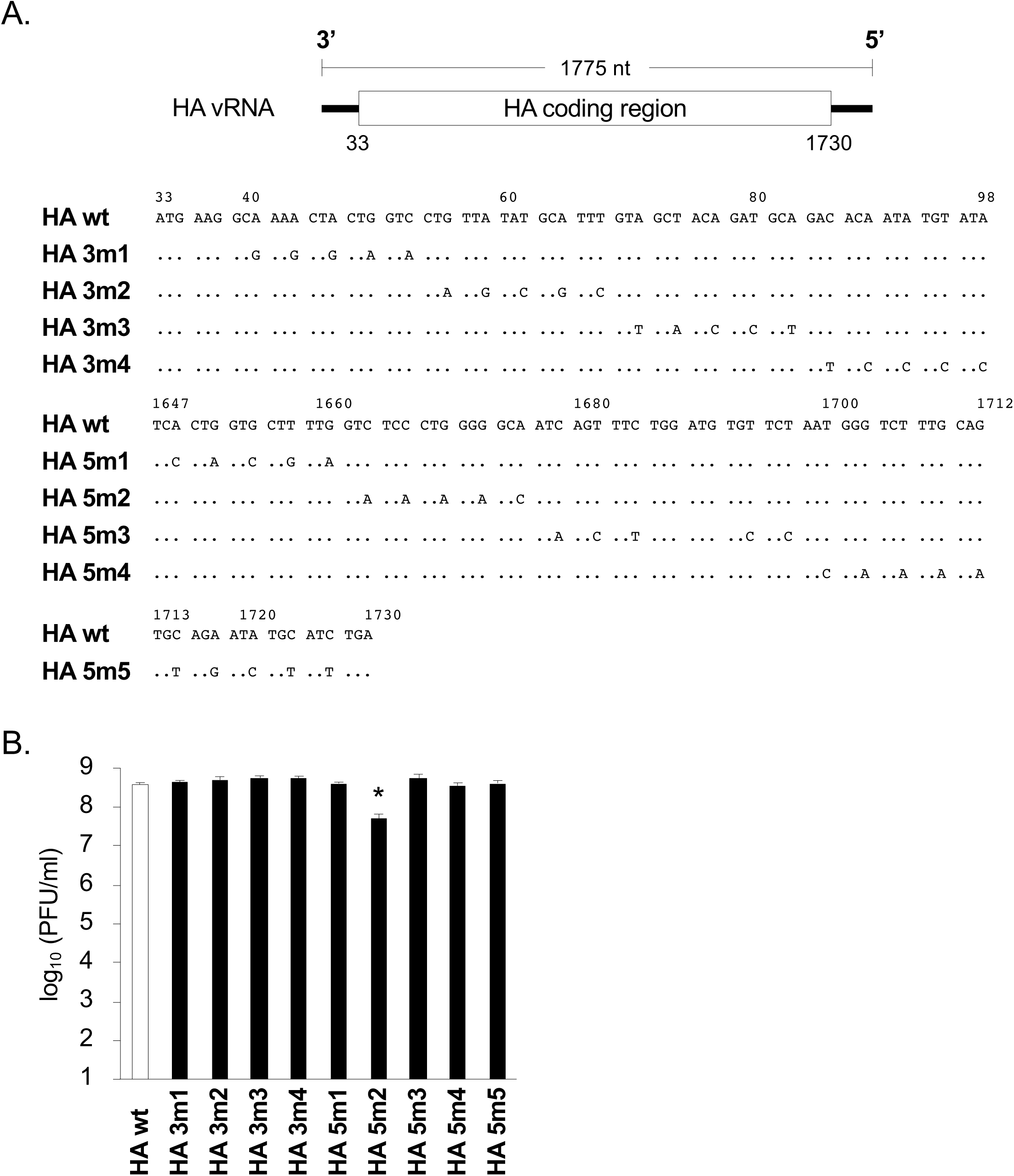
Generation of mutant influenza A viruses by reverse genetics and replication efficiencies of these mutant viruses. (A) Schematic diagram of mutant HA vRNAs with silent mutations introduced into the packaging signal sequences. The 3’ and 5’ ends of the HA coding region (nucleotides 33–98 and 1649–1730, respectively) are shown in the mRNA-sense orientation. (B) 293T cells were transfected with plasmids to produce the mutant and wild-type viruses. MDCK cells were infected with these mutant and wild-type viruses at a multiplicity of infection (MOI) of 10^-5^. Virus yields at 48 h post-infection were determined by a plaque assay on MDCK cells. The data represent the mean ± SD (n = 3). Dunnett’s test; *P < 0.01

### Serial passaging of the HA 5m2 virus in cells

We hypothesized that after several passages, the HA 5m2 virus would acquire adaptive mutations in vRNAs, which would restore viral fitness. Accordingly, the HA 5m2 virus was serially passaged in Madin-Darby Canine Kidney (MDCK) cells, and viral titers were assessed by plaque assay after each passage. The viruses were designated as HA 5m2 P1, P2, P3, P4, P5, P6, P7, P8, P9, and P10 viruses, according to the number of passages in MDCK cells. As expected, the growth of the HA 5m2 P10 virus was restored to approximately 61% of that shown by the wild-type virus (Fig 2A). The titer of the HA 5m2 P10 virus did not increase with an additional 10 passages (data not shown). Sequencing analysis of all eight HA 5m2 P10 virus vRNAs revealed that two mutations were newly introduced into the 5’ terminal coding regions of HA vRNA (T1665C) and PB2 vRNA (G2271T); both of these vRNAs are located within the previously identified genome packaging signaling regions [6, 8, 14, 16].

**Fig 2.**
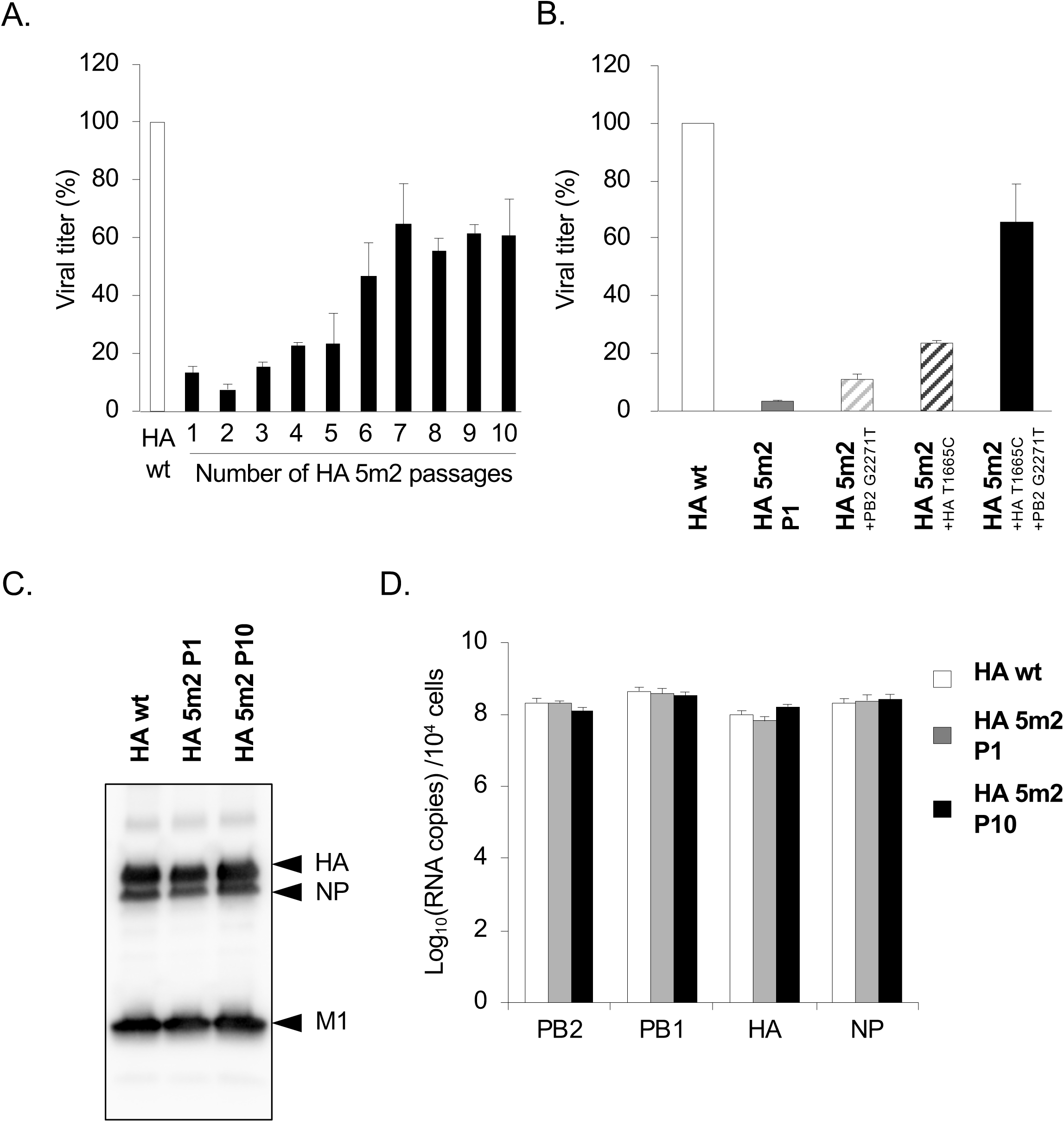
Analysis of HA 5m2 viral replication. (A) The HA 5m2 virus was passaged in MDCK cells at an MOI of 10^-5^. For each passage, supernatants were collected at 48 hours post-infection, and viral yields were determined by a plaque assay on MDCK cells. (B) 293T cells were transfected with plasmids to produce the mutant HA5m2 and wild-type viruses. After infecting MDCK cells at an MOI of 10^-5^, viral yields were determined at 48 hours post-infection. The data represent the mean ± SD (n = 3). (C) The purified viruses were analyzed via western blot using an anti-influenza A virus polyclonal antibody against HA, NP, and M1. (D) MDCK cells were infected with the viruses at an MOI of 1. Total RNA was extracted at 7 hours post-infection. PB2, PB1, HA, and NP vRNAs were analyzed using RT-qPCR analysis. Data represent the mean ±SD of two independent experiments, each performed in triplicate.

To assess whether these two mutations contributed to the growth of the HA 5m2 virus, we used reverse genetics to generate recombinant HA 5m2 viruses with additional mutation(s). Recombinant HA 5m2 viruses, which possessed a single mutation in HA T1665C or PB2 G2271T, both showed replication that was partially restored to 11 and 24% of the wild-type virus, respectively (Fig 2B). The recombinant HA 5m2 virus with a double mutation showed replication that was approximately 65% of the wild-type virus, similar to the replication levels of the HA 5m2 P10 virus (Fig 2A and 2B).

The HA T1665C and PB2 G2271T mutations lead to amino acid substitutions HA S545P and PB2 Q748H, respectively. The amino acid 545 is located in the transmembrane region of the HA protein; hence, this substitution may affect intracellular transport to the plasma membrane and subsequent incorporation of HA proteins into progeny virions. Therefore, we examined the amount of HA protein incorporated into progeny virions. Western blotting showed that the amounts of HA protein were comparable in the HA 5m2 P10, HA 5m2 P1, and wild-type virions. This suggests that the S545P substitution has little or no effect on the amount of HA protein incorporated into virions (Fig 2C). To examine the impact of the Q748H substitution on the polymerase activity of the PB2 protein, we used RT-qPCR to quantify the amount of vRNA in virus-infected cells at 7 hours post-infection. The amount of vRNA was similar in HA 5m2 P10-infected, HA 5m2 P1-infected, and wild-type virus-infected cells, suggesting that the Q748H substitution in PB2 had little or no effect on polymerase activity (Fig 2D). Taken together, these results suggest that T1665C mutations in HA vRNA and G2271T mutations in PB2 vRNA participate in the restoration of HA 5m2 viral replication at the RNA level.

### Efficiency of packaging eight vRNAs in HA 5m2 viruses

Because the HA 5m2 virus possessed five silent mutations in the 5’ packaging signal region of HA vRNA, we predicted that it would show defects in the packaging efficiency of vRNAs (especially of HA vRNA). We also expected that additional mutations in the 5’ packaging signals of HA and PB2 vRNAs would improve the packaging efficiency of HA vRNA. To assess the packaging efficiency of the HA 5m2 P1 and HA 5m2 P10 viruses, we extracted vRNA from wild-type, HA 5m2 P1, and HA 5m2 P10 viruses; then, we quantified the amount of the eight influenza vRNA segments using RT-qPCR. As expected, the HA 5m2 P1 virus showed a marked defect in the packaging efficiency of HA vRNA; the packaging efficiency was reduced to approximately 24% compared with that of the wild-type virus. The HA 5m2 P1 virus also showed small defects in the packaging efficiency of PA, NP, and NA vRNAs (Fig 3A). Importantly, in the HA 5m2 P10 virus, the packaging efficiency of HA vRNA was largely recovered to approximately 72% of that in the wild-type, and those of PA, NP, and NA vRNAs were also partially recovered (Fig 3A).

**Fig 3.**
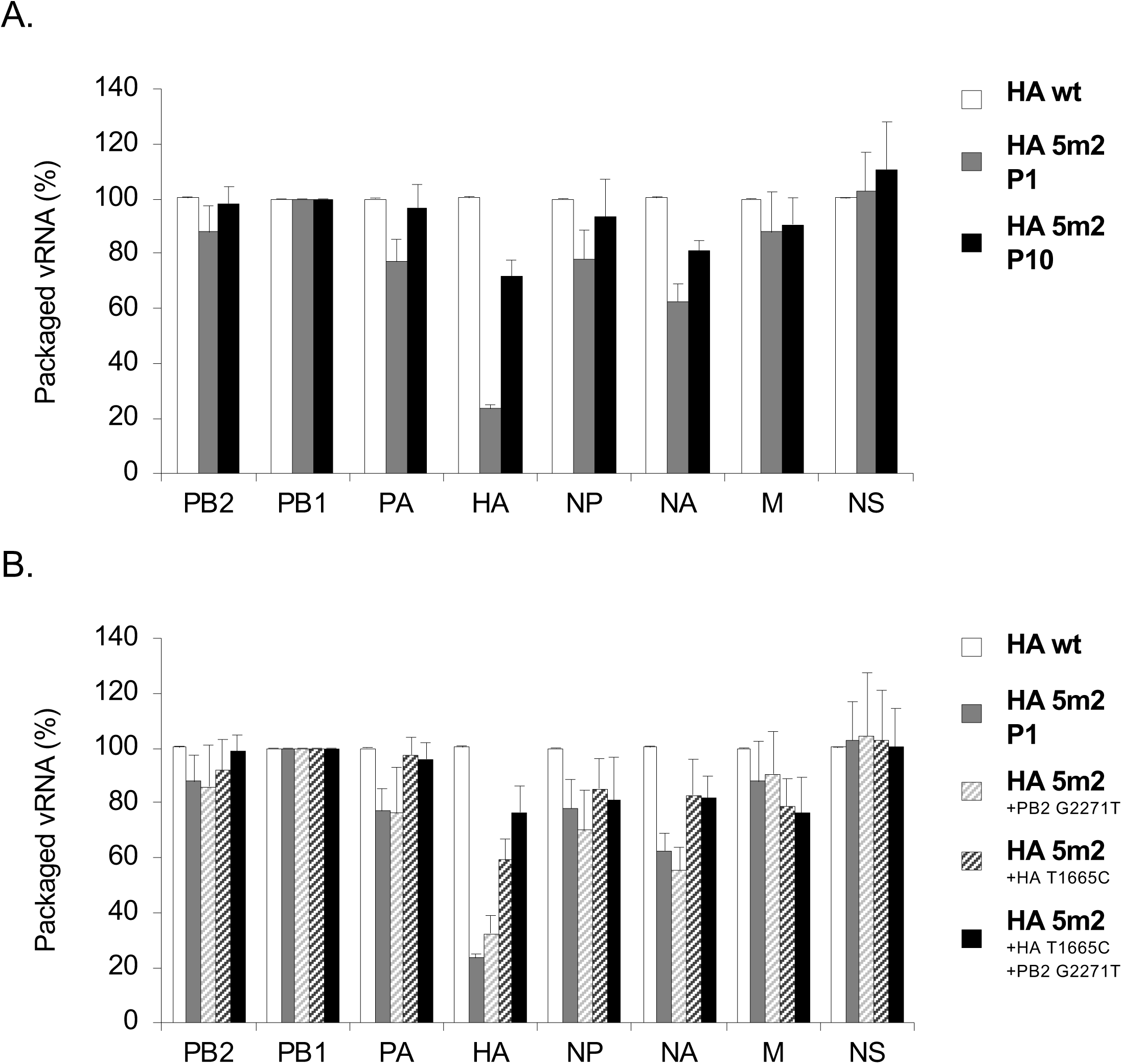
Packaging of individual vRNA segments into progeny viral particles. The amount of vRNA extracted from purified virus particles was quantified by RT-qPCR. All vRNAs were normalized to the amount of PB1 vRNA and to the average of that contained in the wild-type virus. Data represent the mean ±SD from two independent experiments each conducted in triplicate. (A) The panel shows the relative amounts of eight vRNA from the wild-type virus, HA 5m2 P10 virus, and HA 5m2 P1 virus. (B) The panel shows the relative amounts of eight vRNA from the wild-type virus and mutant HA 5m2 viruses.

To further examine the contribution of the PB2 G2271T and HA T1665C mutations to vRNA packaging efficiency, we used RT-qPCR to analyze the amount of packaged vRNA in recombinant HA 5m2 viruses possessing a single mutation or a double mutation. In both recombinant HA 5m2 viruses with either a single PB2 G2271T or HA T1665C mutation, the packaging efficiency of HA vRNA was restored to approximately 33 and 59% to that of the wild-type virus, respectively (Fig 3B). In the recombinant HA 5m2 virus possessing the double mutation, the packaging efficiency of HA vRNA was largely restored to approximately 77% of that of the wild-type virus (Fig 3B); this was consistent with the packaging efficiency of HA vRNA in the HA 5m2 P10 virus. Taken together, these results show that disruption of the 5’ genome packaging signal in HA vRNA reduces the packaging efficiency of HA vRNA. Our results also show that additional mutations in the sequences of 5’ genome packaging signals of HA and PB2 vRNAs are required for efficient HA vRNA packaging. This suggests that functional interactions occur between HA and PB2 vRNAs via their 5’ genome packaging signals during viral replication.

### Ultrastructural analysis of the HA 5m2 viruses

To investigate how the RNPs are packaged into the HA 5m2 P1 and HA 5m2 P10 viruses, we examined ultrathin sections via electron microscopy (EM). Representative images of transversely and longitudinally sectioned wild-type, HA 5m2 P1, and HA 5m2 P10 virions, budding from the cell surfaces, are shown in Fig 4A. While some transversely sectioned wild-type and HA 5m2 P10 virions appeared empty, as reported previously [26], almost all longitudinally sectioned wild-type and HA 5m2 P10 virions contained RNPs positioned at the tip of the budding virions. Conversely, the HA 5m2 P1 viruses often possessed empty particles in both transverse and longitudinal sections, suggesting that the HA 5m2 virus had defects in the incorporation of RNPs.

**Fig 4.**
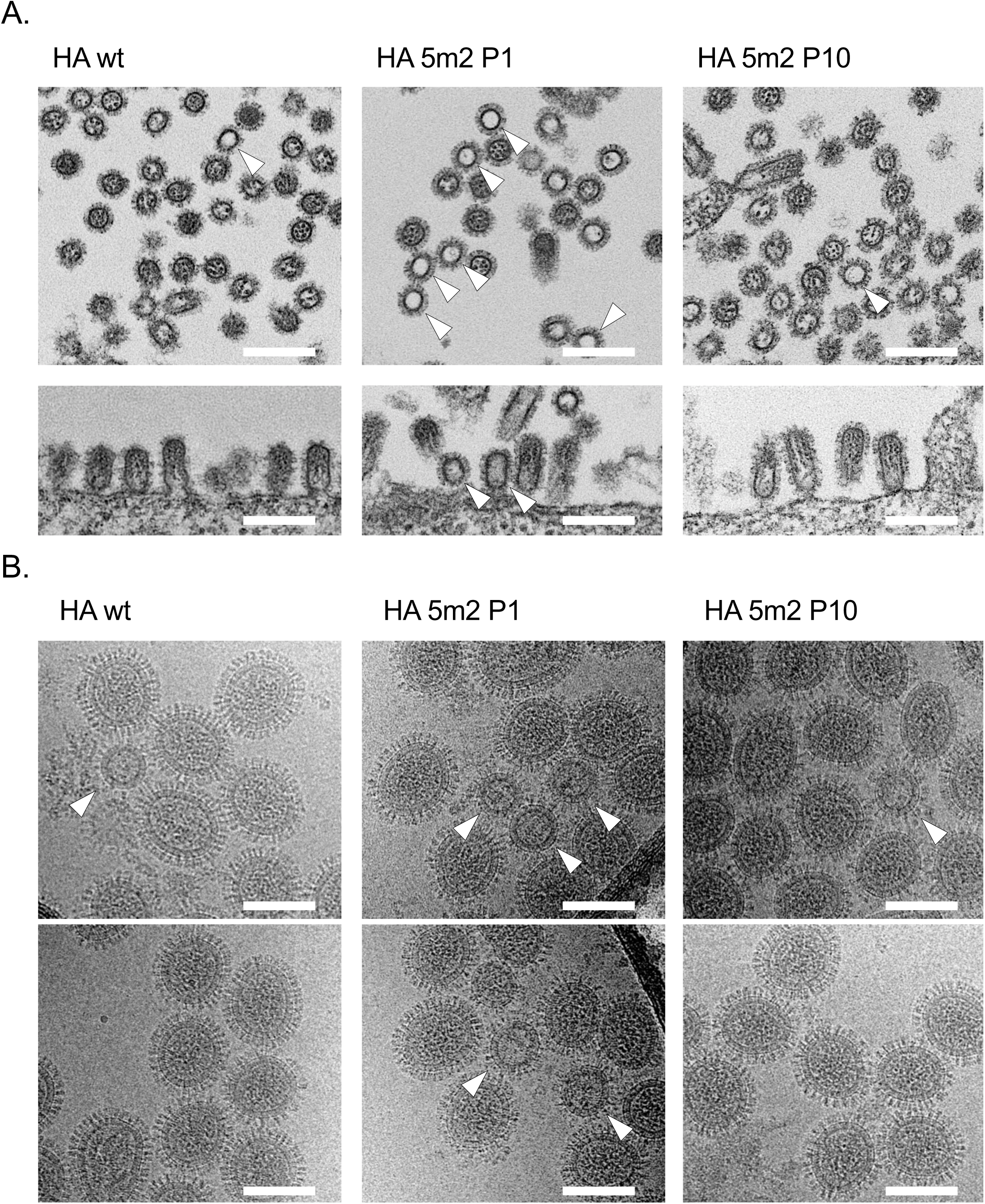
Morphology of HA 5m2 P1, HA5m2 P10, and wild-type viruses. (A) Ultrathin sections of viral particles budding from infected MDCK cells were observed using transverse section (upper side) and longitudinal section (lower side). Empty particles are indicated by arrowheads. Bars, 200 nm. (B) Purified virus particles were observed by cryo-TEM. Empty particles are indicated by arrowheads. Bars, 100 nm.

To examine the incorporation of RNPs into virions in more detail, we determined the proportion of empty particles in wild-type, HA 5m2 P1, and HA 5m2 P10 viruses. For this, virions were purified by ultracentrifugation through a sucrose cushion and observed by cryo-TEM (Fig 4B). The particles of the wild-type virus mainly showed uniformly spherical shapes of approximately 111 nm in diameter; 0.9% (n=1085) of virions appeared empty or contained only a few vRNPs (Table 1). With a diameter of approximately 87 nm (n=10, *p*<0.0001), such vRNP packaging-deficient particles were significantly smaller than intact virions containing multiple RNPs. The proportion of vRNP packaging-deficient particles in the HA 5m2 P1 virus was 7.0% (n=1401), significantly higher than those of the HA 5m2 P10 (3.2%, n=852, *p*<0.0001) and wild-type (0.9%, n=1085, *p*<0.0001) viruses. The results of EM analysis support the notion that potential interactions between the 5’ packaging signals of HA and PB2 vRNAs are important for appropriate genome packaging.

**Table 1.**
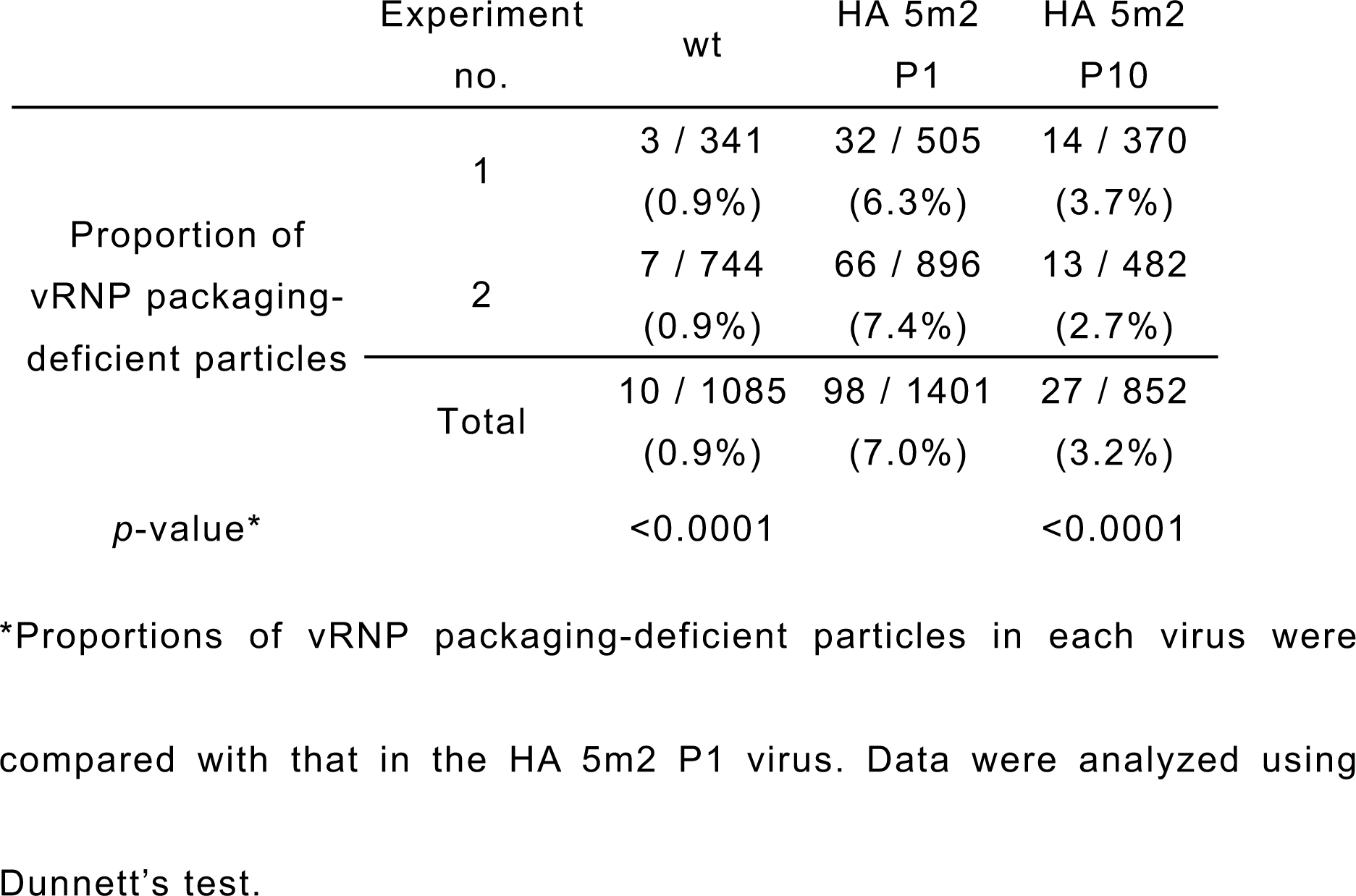
Effect of silent mutations and passages on the packaging of RNPs into progeny virions

### A direct interaction between HA and PB2 vRNA occurs via the 5’ ends of packaging signals *in vitro*

We next aimed to confirm that a functional interaction between the 5’ terminal regions of HA and PB2 vRNAs is involved in efficient packaging of HA vRNA. For this, we used a gel shift assay to examine whether direct RNA-RNA interactions between these two vRNAs occur *in vitro*. To eliminate possible nonspecific interactions via the non-packaging signal regions of vRNAs, we synthesized a short HA vRNA comprising the 5’ noncoding region and the 120-nucleotide long coding region, which is designated as 5’HA(120). We also synthesized a short PB2 vRNA comprising the 5’ noncoding region and the 300-nucleotide long coding region, which is designated as 5’PB2(300) (Fig 5A). In addition to the 5’HA(120) possessing the wild-type HA vRNA sequence, we synthesized mutant 5’HA(120) sequences into which we introduced five silent mutations corresponding to HA 5m1, HA 5m2, HA 5m3, HA 5m4, and HA 5m5 (Fig 1A); these mutant 5’HA(120) vRNAs were designated as 5’HA(120) 5m1, 5’HA(120) 5m2, 5’HA(120) 5m3, 5’HA(120) 5m4, and 5’HA(120) 5m5, respectively. The mixture of wild-type 5’HA(120) and 5’PB2(300) showed slower migration of the band, indicating formation of a vRNA-vRNA complex (Figs 5B and 5C). The mutant 5’HA(120) 5m1, 5’HA(120) 5m3, 5’HA(120) 5m4, and 5’HA(120) 5m5 vRNAs also formed a complex with 5’PB2(300), whose proportions were 69-95% compared to the complex of 5’HA(120) and 5’PB2(300). These results are consistent with our viral replication data, showing that such mutations in HA vRNA did not markedly affect viral growth (Fig 1B). In contrast, the 5’HA(120) 5m2 vRNA associated with 5’PB2(300) to a lesser degree and did not form a vRNA-vRNA complex efficiently, with only 12% complex formation compared to the 5’HA(120) and 5’PB2(300), correlating with the reduced viral growth of the HA 5m2 virus (Fig 1B). Taken together, these results indicate that there is an interaction between the 5’ packaging signals of HA and PB2 vRNAs in the context of vRNPs, which is important for optimal packaging of HA vRNA.

**Fig 5.**
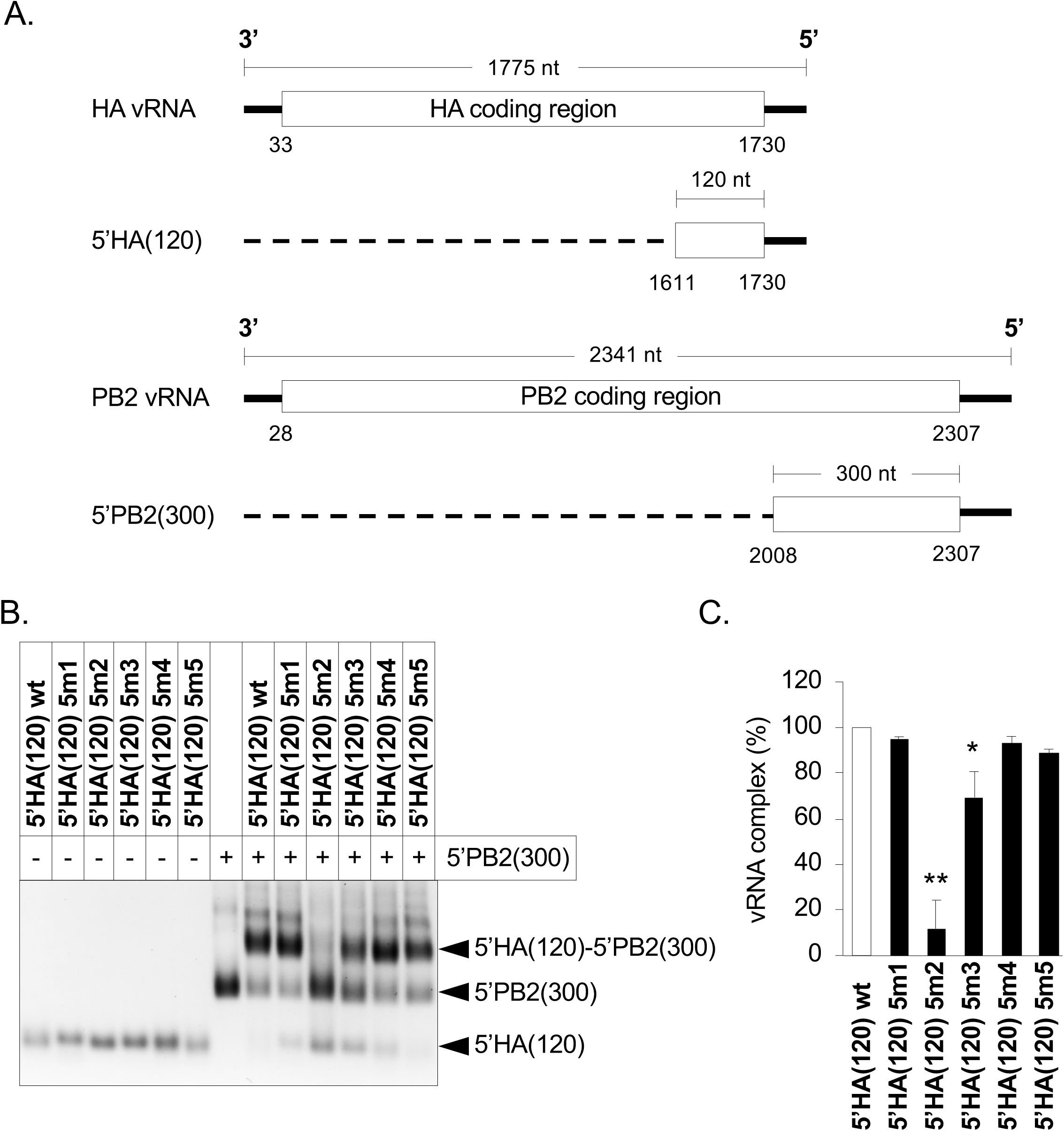
Gel shift assay using 5’ end sequences of HA and PB2 vRNAs. (A) Schematic representation of 5’HA(120) and 5’PB2(300) vRNAs. The solid line represents the noncoding regions of vRNAs at the 3’ and 5’ ends. The white box represents the coding regions of the HA or PB2 proteins. The dashed line represents the deleted regions. 5’HA(120) and 5’PB2(300) contain 120 or 300 nucleotides in the coding and noncoding regions at their 5’ ends, respectively. (B) The effect of silent mutations in 5’HA(120) vRNA on binding to 5’PB2(300) vRNA. 5’PB2(300) vRNA was incubated with wild-type and mutated 5’HA(120) vRNAs as indicated at the top of the gel image. Individual vRNA bands and vRNA complexes are indicated on the right. (C) Quantification of 5’HA(120)-5’ PB2(300) complexes for each lane. The relative band intensity of the complex is indicated in comparison to that of the wild-type. The data represent the mean ± SD (n = 3). Dunnett’s test; *P < 0.01; **P < 0.001.

## Discussion

Specific interactions among eight different vRNAs are likely required for selective genome packaging of the influenza viruses. To identify the interactions of HA vRNA in the context of RNPs, we generated a mutant HA 5m2 virus with reduced packaging efficiency of HA vRNA, and repeatedly passaged this virus in cultured cells to restore viral fitness. We found that the HA 5m2 virus acquired additional mutations in the 5’ packaging signal sequences of HA and PB2 vRNAs; this restored HA vRNA packaging efficiency and viral growth. Our results suggest that the packaging signal at the 5’ terminal coding region of HA vRNA is involved in co-packaging of the eight different vRNAs; this likely occurs via a direct RNA-RNA interaction with the 5’ packaging signal of PB2 vRNA. To the best of our knowledge, this is the first study experimentally showing that a packaging-deficient virus can recover its packaging efficiency by the introduction of mutations in other vRNA segments during viral replication.

The 5’ packaging signal sequence, located at nucleotides 1662 to 1681 in HA vRNA, is more than 90% conserved in H1 subtype influenza viruses. However, the neighboring regions show less sequence conservation [14], suggesting that this region is important for HA vRNA packaging at the RNA level. Our results show that introduction of five silent mutations into the highly conserved region at nucleotides 1664 to 1676 in HA vRNA reduced the incorporation efficiency of HA vRNA (Fig. 2A). An additional mutation at nucleotide 1665 restored the reduced efficiency of HA vRNA packaging (Fig. 2B). This confirms that the highly conserved region in the sequence of the 5’ terminal packaging signal is involved in incorporation of HA vRNA. Recent findings indicate that NP non-uniformly decorates vRNA [1, 2]. Therefore, it is possible that the region at nucleotides 1664 to 1676 in HA vRNA, identified in this study, is NP-free and forms secondary or tertiary structures on vRNPs to interact with the 5’ packaging signal of PB2 vRNA. However, whether the 5’ terminal coding region of HA vRNA is a low NP-binding region remains unclear [1, 2]. Additional work is necessary to determine the precise location of the NP-free region of HA vRNA in the A/WSN/33 strain.

After the HA 5m2 virus was serially passaged in MDCK cells, the virus acquired a G2271T mutation in the 5’ packaging signal sequence of PB2 vRNA, which recovered the reduced incorporation of HA vRNA (Fig. 2B). This finding suggests that the region around nucleotide 2271 in PB2 vRNA is involved in interactions with HA vRNA. However, a previous study showed that the introduction of silent mutations at nucleotides 2268 to 2286, including a mutation at nucleotide 2271, in PB2 vRNA did not reduce the packaging efficiency of HA vRNA [14]. Therefore, it is possible that the mutation at G2271T in PB2 vRNA, found in this study, was acquired for the optimal packaging of HA 5m2 vRNA but not of wild-type HA vRNA.

Although the 5’ terminal coding region of H1 subtype HA vRNA is conserved in H1 HA vRNA, it is substantially different from other subtypes of HA vRNAs at the nucleotide level [13]. In contrast, the 5’ terminal coding region of PB2 vRNA is highly conserved in influenza viruses [13, 17], suggesting that intersegmental interactions between HA and PB2 vRNAs via their 5’ terminal regions may be specific to the H1 subtype. Native gel electrophoresis analysis of *in-vitro* vRNA-vRNA interactions revealed that avian H5N2 and human H3N2 viruses show different intersegmental networks among the eight vRNAs, and do not show strong interactions between HA and PB2 vRNAs [22, 24]. In the context of vRNPs, *in-vitro* interactions do not necessarily reflect interactions *in-vivo*. Therefore, further studies are needed to clarify whether HA vRNAs of other subtypes require interactions with PB2 vRNA for HA vRNA packaging in the context of vRNPs.

The reduced HA vRNA packaging efficiency of the HA 5m2 virus was rescued when the virus spontaneously acquired adaptive mutations in HA and PB2 vRNAs during serial passaging (Fig. 2A and 2B). Previously, Marsh et al., serially passaged an influenza A virus (WSN strain), which possesses synonymous mutations in HA vRNAs, to determine whether the virus would generate adaptive mutations for recovery of the reduced HA vRNA packaging efficiency. However, the virus did not generate any mutations to improve viral fitness [14]. This may have been due to the number of synonymous mutations introduced into the HA vRNAs. Nine nucleotide mutations were introduced into the region of 1659 to 1673 in the HA vRNAs in the study by Marsh et al., while only five mutations were introduced into the region of 1664 to 1679 in HA vRNAs in our present study. More mutations may cause severe incompatibility in vRNA-vRNA interactions. Even repeated serial passaging may not generate multiple nucleotide changes in HA vRNA and its respective interacting vRNA to restore the HA vRNA packaging efficiency.

In conclusion, we have shown that an interaction between HA vRNA and PB2 vRNA via the 5’ packaging signals is important for HA vRNA packaging. Our results suggest that HA vRNA is co-packaged with PB2 vRNA into virions. These findings will help us understand how reassortant influenza viruses incorporate HA vRNA segments, and how pandemic viruses emerge via genetic reassortment.

## Materials and Methods

### Cells

Human embryonic kidney (293T) cells were maintained in Dulbecco’s Modified Eagle Medium (D6046, Sigma) supplemented with 10% fetal bovine serum (FB-1365, Biosera, Chile). Madin-Darby Canine Kidney (MDCK) cells were grown in Minimal Essential Medium (MEM)(11430-030, Gibco) containing 5% newborn calf serum (16010-159, Gibco, New Zealand). Cultures were maintained at 37°C in a 5% CO2 atmosphere. The medium used during viral infection of cells was MEM containing 0.3% bovine serum albumin (BSA/MEM).

### Construction of the PolI HA plasmid

To generate the PolI HA mutant plasmid, we amplified the PolI-HA plasmid by inverse PCR using a previously published protocol [27]. The primers used are listed in S1 Table.

### Reverse genetics

Reverse genetics were performed using PolI plasmids that contained cDNA sequences of the A/WSN/33(H1N1) viral genes; all procedures were conducted as described previously [28]. Briefly, eight PolI plasmids and pCAGGS protein-expression plasmids for PB2, PB1, PA, and NP were mixed with the transfection reagent TransIT-293 (Mirus), and added to 293T cells cultured in BSA/MEM. At 48 hours post-transfection, the cells were treated with 1 μg /ml of TPCK-Trypsin for 30 min, centrifuged at 1750 × g for 15 min at 4°C, and the supernatant was harvested and stored at −80°C. To generate mutant viruses, the PolI-HA wt plasmid was replaced with a PolI-HA mutant plasmid. Viral titers were determined by a plaque assay conducted using MDCK cells.

### Virus purification

After collecting the supernatants from virus-infected MDCK cells, each supernatant was clarified by centrifugation at 1750 × g for 15 min at 4°C, followed by another centrifugation at 6700 × g for 30 min at 4°C. To eliminate RNAs outside of the virus particles, the supernatants were treated with 5 μg/ml RNase A (Nacalai Tesque) for 1 hour at 37°C. Virions in the supernatants were purified by ultracentrifugation through a 30% (w/w) sucrose cushion at 125,000 × g for 2 hours at 4°C. The pellets were then resuspended in phosphate buffered saline (PBS).

### Western blotting

MDCK cells were infected with the viruses at a multiplicity of infection (MOI) of 1 on ice for 1 hour. The infected MDCK cells were then cultured in BSA/MEM containing 1 μg /ml TPCK-Trypsin. At 10 hours post-infection, the supernatant was harvested and purified as described above. The purified virus was dissolved with an equal volume of 2x Tris-Glycine SDS Sample Buffer (Thermo Fisher Scientific) and boiled for 5 min without a reducing agent, and then subjected to SDS-PAGE. Proteins were electroblotted onto Immobilon-P transfer membranes (Millipore Corporation). The membranes were blocked with Blocking One (Nacalai Tesque) for 30 min at room temperature, and then incubated with goat anti-influenza A virus polyclonal antibody (ab20841, Abcam, 1:10,000 dilution) overnight at 4°C. After incubation with rabbit anti-goat IgG secondary antibody (ab6741, Abcam, 1:10,000 dilution) for 1 hour at room temperature, the blots were developed using a Chemi-Lumi One Super (Nacalai Tesque).

### RT-qPCR

The packaged vRNAs were extracted from purified viruses using an RNeasy Mini Kit (Qiagen). 100 ng of extracted vRNAs were reverse transcribed using a Uni-12 primer (5’-AGCRAAAGCAGG-3’) and Superscript III reverse transcriptase (Invitrogen). Quantification was performed by qPCR on a Rotor-Gene Q 2plex System (Qiagen) using segment-specific primers modified from the protocol by Marsh *et al*. The primers used are listed in S1 Table. For each sample, reactions contained 1 μl 10-fold diluted RT product, 7.5 μl THUNDERBIRD SYBR qPCR Mix (Toyobo), 0.25 μM segment-specific primers, for a final volume of 15 μl. Cycling conditions were: 2 min at 94°C, followed by 40 cycles (98°C for 10 s, 55°C for 15 s, and 72°C for 30 s).

In Figure 2D, total RNA was extracted from virus-infected cells using an RNeasy Mini Kit (Qiagen). For PB2, PB1, HA, and NP vRNAs, RT-qPCR was performed as described above, using 100 ng total RNA. The values were expressed as numbers of RNA copies in an infected cell, assuming that a cell contained 10 pg of RNA [29, 30].

### Ultrathin section electron microscopy (EM)

Ultrathin section EM was performed as described previously [3]. MDCK cells were infected at MOI=10. At 10 hours post-infection, the infected cells were prefixed with 2.5% glutaraldehyde in 0.1 M cacodylate buffer (pH 7.4) on ice. Ultrathin (50-nm-thick) sections were stained with 2% uranyl acetate in 70% ethanol and in Reynold’s lead citrate. Representative images were acquired using HT7700 (Hitachi).

### Quantification of vRNP packaging-deficient particles

Purified viruses were applied onto C-flat holey carbon grids (Protichips), which were blotted on a Vitrobot Mark IV (Thermo Fisher Scientific) before plunge-freezing in liquid ethane. Samples were kept cool in liquid nitrogen. Samples were imaged at 200 kV on a Talos F200C (Thermo Fisher Scientific) equipped with a Ceta 16M CMOS camera (Gatan). More than 300 particles were observed for each experiment.

### *In vitro* RNA synthesis

Shortened HA and PB2 vRNAs were synthesized *in vitro* using T7 transcription as described previously [30]. Briefly, templates containing a T7 phage promoter sequence (5’-TAATACGACTCACTATAGGG-3’) were amplified by PCR using corresponding primer pairs and were purified with a QIAquick PCR purification kit (Qiagen). The primers used are listed in S1 Table. Purified PCR products were transcribed *in vitro* using the RiboMAX Large Scale RNA Production System-T7 (Promega) at 37°C for 4 h, followed by RQ1 DNase I (Promega) digestion of the DNA template at 37°C for 15 min. The transcript was purified with an RNeasy Mini Kit.

### RNA binding assay

RNA-RNA interactions were analyzed by electrophoretic mobility shift assays essentially as described previously [24]. To facilitate RNA folding, adjustments were made to the reaction buffer employed and incubation time as described below [31]. Pairs of purified vRNAs (2 pmol of each vRNA) were denatured for 10 min at 65°C in 5 µl of ultrapure water and cooled on ice. We then added 5 μl of 2-fold concentrated buffer (final concentration: 50 mM HEPES, 50 mM KCl, and 20 mM MgCl2) and incubated the samples for 2 hours at 37°C. Then, 2 µl of loading buffer [40% (v/v) glycerol and 0.05% (w/v) bromophenol blue] was added to the samples, and the samples were analyzed using 1.0% agarose gels containing 0.01% (w/v) ethidium bromide. Native gel electrophoresis of the RNA complexes was performed at 4°C in a buffer containing 50 mM Tris, 44.5 mM borate, and 0.1 mM MgCl2. We then determined the RNA weight fraction (%) of each band in each lane. The percentage of intermolecular complex formation was determined by dividing the weight fraction of a band by the sum of weight fractions in the corresponding lane.

### Statistical analysis

The statistical significance of the viral titers, the proportions of vRNP packaging-deficient particles and the formation efficiencies of vRNA-vRNA complexes were calculated using Dunnett’s test. Diameters of virus particles observed by EM were statistically analyzed using Welch’s t-test. P values of < 0.01 were considered significant.

## Acknowledgments

We thank Connor Park for editing the manuscript.

## References

1. Williams GD, Townsend D, Wylie KM, Kim PJ, Amarasinghe GK, Kutluay SB, et al. Nucleotide resolution mapping of influenza A virus nucleoprotein-RNA interactions reveals RNA features required for replication. Nature communications. 2018;9(1):465. Epub 2018/02/02. doi: 10.1038/s41467-018-02886-w. PubMed PMID: 29386621; PubMed Central PMCID: PMCPMC5792457.

2. Lee N, Le Sage V, Nanni AV, Snyder DJ, Cooper VS, Lakdawala SS. Genome-wide analysis of influenza viral RNA and nucleoprotein association. Nucleic acids research. 2017;45(15):8968–77. doi: 10.1093/nar/gkx584.

3. Noda T, Sagara H, Yen A, Takada A, Kida H, Cheng RH, et al. Architecture of ribonucleoprotein complexes in influenza A virus particles. Nature. 2006;439(7075):490–2. Epub 2006/01/27. doi: 10.1038/nature04378. PubMed PMID: 16437116.

4. Chou YY, Vafabakhsh R, Doganay S, Gao Q, Ha T, Palese P. One influenza virus particle packages eight unique viral RNAs as shown by FISH analysis. Proceedings of the National Academy of Sciences of the United States of America. 2012;109(23):9101–6. Epub 2012/05/02. doi: 10.1073/pnas.1206069109. PubMed PMID: 22547828; PubMed Central PMCID: PMCPMC3384215.

5. Fujii Y, Goto H, Watanabe T, Yoshida T, Kawaoka Y. Selective incorporation of influenza virus RNA segments into virions. Proceedings of the National Academy of Sciences of the United States of America. 2003;100(4):2002–7. Epub 2003/02/08. doi: 10.1073/pnas.0437772100. PubMed PMID: 12574509; PubMed Central PMCID: PMCPMC149948.

6. Watanabe T, Watanabe S, Noda T, Fujii Y, Kawaoka Y. Exploitation of nucleic acid packaging signals to generate a novel influenza virus-based vector stably expressing two foreign genes. Journal of virology. 2003;77(19):10575–83. Epub 2003/09/13. PubMed PMID: 12970442; PubMed Central PMCID: PMCPMC228515.

7. Fujii K, Fujii Y, Noda T, Muramoto Y, Watanabe T, Takada A, et al. Importance of both the coding and the segment-specific noncoding regions of the influenza A virus NS segment for its efficient incorporation into virions. Journal of virology. 2005;79(6):3766–74. Epub 2005/02/26. doi: 10.1128/jvi.79.6.3766-3774.2005. PubMed PMID: 15731270; PubMed Central PMCID: PMCPMC1075679.

8. Muramoto Y, Takada A, Fujii K, Noda T, Iwatsuki-Horimoto K, Watanabe S, et al. Hierarchy among viral RNA (vRNA) segments in their role in vRNA incorporation into influenza A virions. Journal of virology. 2006;80(5):2318–25. Epub 2006/02/14. doi: 10.1128/jvi.80.5.2318-2325.2006. PubMed PMID: 16474138; PubMed Central PMCID: PMCPMC1395381.

9. Ozawa M, Fujii K, Muramoto Y, Yamada S, Yamayoshi S, Takada A, et al. Contributions of two nuclear localization signals of influenza A virus nucleoprotein to viral replication. Journal of virology. 2007;81(1):30–41. Epub 2006/10/20. doi: 10.1128/jvi.01434-06. PubMed PMID: 17050598; PubMed Central PMCID: PMCPMC1797272.

10. Ozawa M, Maeda J, Iwatsuki-Horimoto K, Watanabe S, Goto H, Horimoto T, et al. Nucleotide sequence requirements at the 5’ end of the influenza A virus M RNA segment for efficient virus replication. Journal of virology. 2009;83(7):3384–8. Epub 2009/01/23. doi: 10.1128/jvi.02513-08. PubMed PMID: 19158245; PubMed Central PMCID: PMCPMC2655591.

11. Goto H, Muramoto Y, Noda T, Kawaoka Y. The genome-packaging signal of the influenza A virus genome comprises a genome incorporation signal and a genome-bundling signal. Journal of virology. 2013;87(21):11316–22. Epub 2013/08/09. doi: 10.1128/jvi.01301-13. PubMed PMID: 23926345; PubMed Central PMCID: PMCPMC3807325.

12. Liang Y, Hong Y, Parslow TG. cis-Acting packaging signals in the influenza virus PB1, PB2, and PA genomic RNA segments. Journal of virology. 2005;79(16):10348–55. Epub 2005/07/30. doi: 10.1128/jvi.79.16.10348-10355.2005. PubMed PMID: 16051827; PubMed Central PMCID: PMCPMC1182667.

13. Gog JR, Afonso Edos S, Dalton RM, Leclercq I, Tiley L, Elton D, et al. Codon conservation in the influenza A virus genome defines RNA packaging signals. Nucleic acids research. 2007;35(6):1897–907. Epub 2007/03/03. doi: 10.1093/nar/gkm087. PubMed PMID: 17332012; PubMed Central PMCID: PMCPMC1874621.

14. Marsh GA, Hatami R, Palese P. Specific residues of the influenza A virus hemagglutinin viral RNA are important for efficient packaging into budding virions. Journal of virology. 2007;81(18):9727–36. Epub 2007/07/20. doi: 10.1128/jvi.01144-07. PubMed PMID: 17634232; PubMed Central PMCID: PMCPMC2045411.

15. Hutchinson EC, Curran MD, Read EK, Gog JR, Digard P. Mutational analysis of cis-acting RNA signals in segment 7 of influenza A virus. Journal of virology. 2008;82(23):11869–79. Epub 2008/09/26. doi: 10.1128/jvi.01634-08. PubMed PMID: 18815307; PubMed Central PMCID: PMCPMC2583641.

16. Liang Y, Huang T, Ly H, Parslow TG, Liang Y. Mutational analyses of packaging signals in influenza virus PA, PB1, and PB2 genomic RNA segments. Journal of virology. 2008;82(1):229–36. Epub 2007/10/26. doi: 10.1128/jvi.01541-07. PubMed PMID: 17959657; PubMed Central PMCID: PMCPMC2224372.

17. Marsh GA, Rabadan R, Levine AJ, Palese P. Highly conserved regions of influenza a virus polymerase gene segments are critical for efficient viral RNA packaging. Journal of virology. 2008;82(5):2295–304. Epub 2007/12/21. doi: 10.1128/jvi.02267-07. PubMed PMID: 18094182; PubMed Central PMCID: PMCPMC2258914.

18. Hutchinson EC, Wise HM, Kudryavtseva K, Curran MD, Digard P. Characterisation of influenza A viruses with mutations in segment 5 packaging signals. Vaccine. 2009;27(45):6270–5. Epub 2009/10/21. doi: 10.1016/j.vaccine.2009.05.053. PubMed PMID: 19840659; PubMed Central PMCID: PMCPMC2771075.

19. Gao Q, Chou YY, Doganay S, Vafabakhsh R, Ha T, Palese P. The influenza A virus PB2, PA, NP, and M segments play a pivotal role during genome packaging. Journal of virology. 2012;86(13):7043–51. Epub 2012/04/26. doi: 10.1128/jvi.00662-12. PubMed PMID: 22532680; PubMed Central PMCID: PMCPMC3416350.

20. Chou YY, Heaton NS, Gao Q, Palese P, Singer RH, Lionnet T. Colocalization of different influenza viral RNA segments in the cytoplasm before viral budding as shown by single-molecule sensitivity FISH analysis. PLoS pathogens. 2013;9(5):e1003358. Epub 2013/05/15. doi: 10.1371/journal.ppat.1003358. PubMed PMID: 23671419; PubMed Central PMCID: PMCPMC3649991.

21. Lakdawala SS, Wu Y, Wawrzusin P, Kabat J, Broadbent AJ, Lamirande EW, et al. Influenza a virus assembly intermediates fuse in the cytoplasm. PLoS pathogens. 2014;10(3):e1003971. Epub 2014/03/08. doi: 10.1371/journal.ppat.1003971. PubMed PMID: 24603687; PubMed Central PMCID: PMCPMC3946384.

22. Fournier E, Moules V, Essere B, Paillart JC, Sirbat JD, Isel C, et al. A supramolecular assembly formed by influenza A virus genomic RNA segments. Nucleic acids research. 2012;40(5):2197–209. Epub 2011/11/15. doi: 10.1093/nar/gkr985. PubMed PMID: 22075989; PubMed Central PMCID: PMCPMC3300030.

23. Noda T, Sugita Y, Aoyama K, Hirase A, Kawakami E, Miyazawa A, et al. Three-dimensional analysis of ribonucleoprotein complexes in influenza A virus. Nature communications. 2012;3:639. Epub 2012/01/26. doi: 10.1038/ncomms1647. PubMed PMID: 22273677; PubMed Central PMCID: PMCPMC3272569.

24. Gavazzi C, Isel C, Fournier E, Moules V, Cavalier A, Thomas D, et al. An in vitro network of intermolecular interactions between viral RNA segments of an avian H5N2 influenza A virus: comparison with a human H3N2 virus. Nucleic acids research. 2013;41(2):1241–54. Epub 2012/12/12. doi: 10.1093/nar/gks1181. PubMed PMID: 23221636; PubMed Central PMCID: PMCPMC3553942.

25. Gilbertson B, Zheng T, Gerber M, Printz-Schweigert A, Ong C, Marquet R, et al. Influenza NA and PB1 Gene Segments Interact during the Formation of Viral Progeny: Localization of the Binding Region within the PB1 Gene. Viruses. 2016;8(8). Epub 2016/08/25. doi: 10.3390/v8080238. PubMed PMID: 27556479; PubMed Central PMCID: PMCPMC4997600.

26. Gavazzi C, Yver M, Isel C, Smyth RP, Rosa-Calatrava M, Lina B, et al. A functional sequence-specific interaction between influenza A virus genomic RNA segments. Proceedings of the National Academy of Sciences of the United States of America. 2013;110(41):16604–9. Epub 2013/09/27. doi: 10.1073/pnas.1314419110. PubMed PMID: 24067651; PubMed Central PMCID: PMCPMC3799358.

27. Ochman H, Gerber AS, Hartl DL. Genetic applications of an inverse polymerase chain reaction. Genetics. 1988;120(3):621–3. Epub 1988/11/01. PubMed PMID: 2852134; PubMed Central PMCID: PMCPMC1203539.

28. Neumann G, Watanabe T, Ito H, Watanabe S, Goto H, Gao P, et al. Generation of influenza A viruses entirely from cloned cDNAs. Proceedings of the National Academy of Sciences of the United States of America. 1999;96(16):9345–50. Epub 1999/08/04. PubMed PMID: 10430945; PubMed Central PMCID: PMCPMC17785.

29. Hatada E, Hasegawa M, Mukaigawa J, Shimizu K, Fukuda R. Control of Influenza Virus Gene Expression: Quantitative Analysis of Each Viral RNA Species in Infected Cells. The Journal of Biochemistry. 1989;105(4):537–46.

30. Kawakami E, Watanabe T, Fujii K, Goto H, Watanabe S, Noda T, et al. Strand-specific real-time RT-PCR for distinguishing influenza vRNA, cRNA, and mRNA. Journal of virological methods. 2011;173(1):1–6. Epub 2010/12/28. doi: 10.1016/j.jviromet.2010.12.014. PubMed PMID: 21185869; PubMed Central PMCID: PMCPMC3049850.

31. Buchmueller KL, Weeks KM. Tris-borate is a poor counterion for RNA: a cautionary tale for RNA folding studies. Nucleic acids research. 2004;32(22):e184. Epub 2004/12/17. doi: 10.1093/nar/gnh182. PubMed PMID: 15601995; PubMed Central PMCID: PMCPMC545480.

